# Uman Type NF-L Antibodies Are Effective Reagents for the Imaging of Neurodegeneration

**DOI:** 10.1101/2022.08.27.504533

**Authors:** Gerry Shaw, Irina Madorsky, Ying Li, YongSheng Wang, Sabhya Rana, David D. Fuller

## Abstract

Recent work shows that certain immunological assays for the neurofilament light chain NF-L detect informative signals in the CSF and blood of human and animals affected by a variety of CNS injury and disease states. Much of this work has been performed using two mouse monoclonal antibodies to NF-L, UD1 and UD2, also known as 2.1 and 47.3 respectively. These are the essential components of the Uman Diagnostics NF-Light^™^ ELISA kit, the Quanterix Simoa^™^ bead based NF-L assay and others. We show here that the antibodies bind to neighboring epitopes in a short, conserved and unusual peptide in the NF-L “rod” Coil 2 region. We also describe a surprising and useful feature of Uman and similar reagents. While other well characterized NF-L antibodies show robust staining of countless cells and processes in CNS sections from healthy rats, both Uman antibodies reveal only a minor subset of presumably spontaneously degenerating or degenerated neurons and their processes. However following experimental mid-cervical injuries to rat spinal cord both Uman antibodies recognize numerous profiles in tissue sections. The Uman positive material was associated with fiber tracts expected to be damaged by the injury administered and the profiles had the swollen, beaded, discontinuous and sinusoidal morphology expected for degenerating and degenerated processes. We also found that several antibodies to the C terminal “tail” region of NF-L stain undamaged axonal profiles but fail to recognize the Uman positive material. The unmasking of the Uman epitopes and the loss of the NF-L tail epitopes can be mimicked by treating sections from healthy animals with proteases suggesting that the immunological changes we have discovered are due to neurodegeneration induced proteolysis. We have also generated a novel panel of monoclonal and polyclonal antibody reagents directed against the region of NF-L including the Uman epitopes which have staining properties identical to the Uman reagents. Using these we show that the NF-L region to which the Uman reagents bind contains further hidden epitopes distinct from those recognized by the two Uman reagents. We speculate that the Uman type epitopes are part of a binding region important for higher order neurofilament assembly. The work provides important insights into the properties of the NF-L biomarker, describes novel and useful properties of Uman type and NF-L tail binding antibodies and provides a hypothesis relevant to further understanding of neurofilament assembly.

## Introduction

A major focus of recent research has been the identification of potential CNS damage and disease biomarkers used diagnostically, prognostically or for monitoring responses to treatment. Axons are particularly sensitive to CNS damage and disease^1–3^ and packed with neurofilaments (NF). Quantitation of the levels of NF subunit proteins or their protein fragments in CSF and blood is therefore likely to usefully reflect ongoing axonal loss. Previously we described the detection of the phosphorylated axonal form of the NF heavy chain (pNF-H) as a viable blood biomarker of axonal damage and degeneration.^4,5^ The neurofilament light chain NF-L, also known as Nfl or by the HGNC name NEFL, is, like pNF-H, highly abundant, neuron specific and heavily concentrated in axons. Recently numerous studies have focused on NF-L as a biomarker of axonal loss which can be sensitively detected in human and animal CSF and blood.^6–14^

NF-L and pNF-H are members of the intermediate or 10nm filament family consisting of over 60 human gene products including the NF subunits NF-M, α-internexin and peripherin as well as GFAP, vimentin, epithelial keratins and nuclear lamins. All are based on a tripartite structure deduced from their amino acid sequence^15^ now confirmed by structural studies.^16^ An N-terminal globular “head” region is followed by a central α-helical “rod” region with two short non-helical linkers followed by a C-terminal globular “tail” segment (**Fig. 1A**). The α-helical regions have the hydrophobic heptad repeats:^17^ Sets of seven amino acids can be labeled from “a” to “g” repetitively down the amino acid sequence and in the correct register the a and d positions are typically hydrophobic. In the case of NF-L and all other 10nm filament subunits each α-helical region forms a stable parallel dimer with another similar polypeptide due to the hydrophobic a and d amino acids in one α-helix interacting with those in another similar α-helix, allowing the two α-helices to wrap around each other forming an α-helical coiled coil.^15^ Amino acids at the b, c, e, f and g positions are predominantly charged and are presented external to the core of the coiled coil structure. The Coil 2 region contains two conserved “stutters” in which the heptad repeat pattern breaks down, which can be regarded as the insertion of 4 extra amino acids into the heptad pattern.^18^ Coiled coil dimers associate laterally and end to end with other similar dimers to produce tetramers and higher order structures, a series of interactions which will finally produce an assembled 10nm diameter NF.^19^

**Figure 1:**
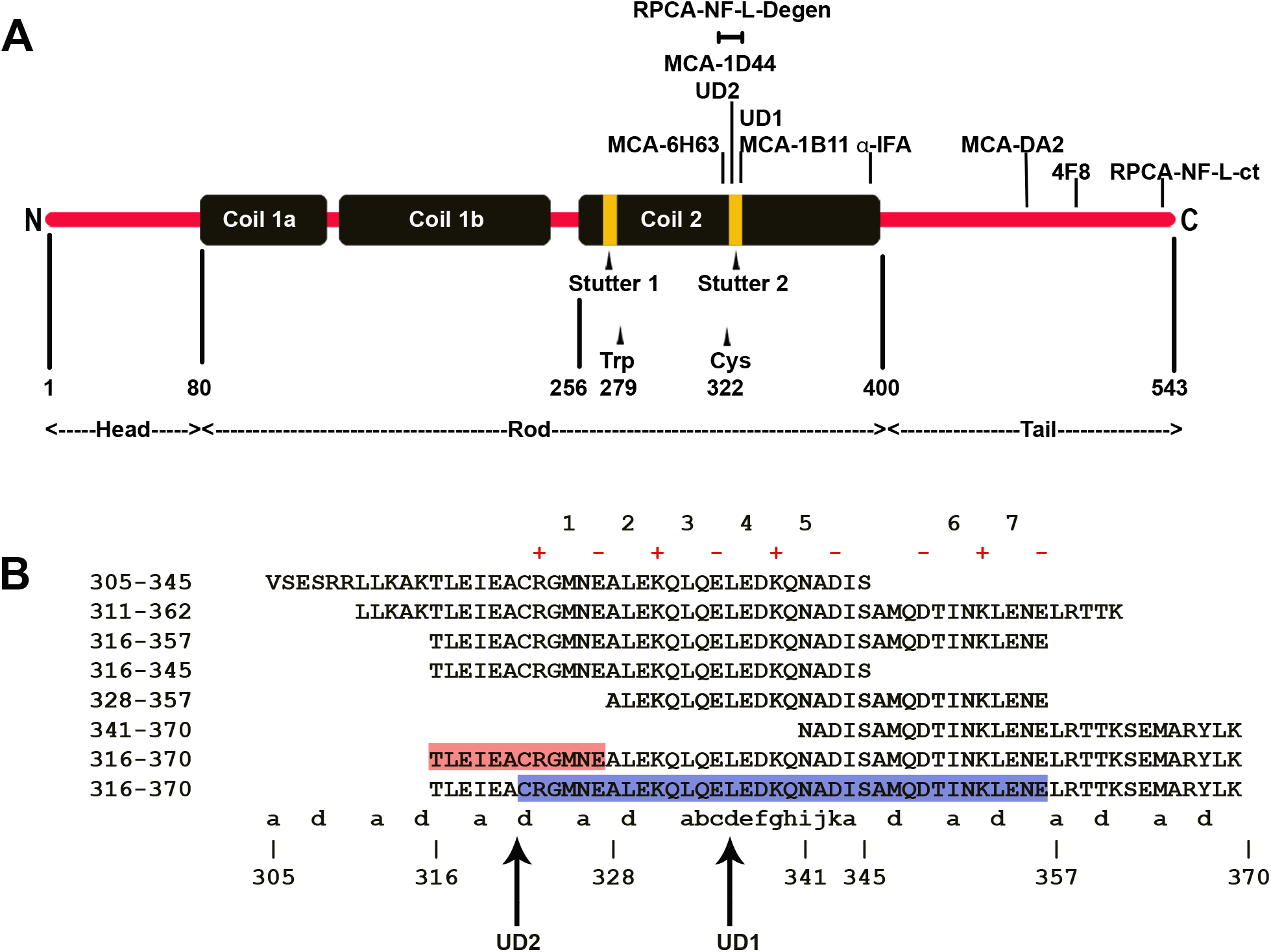
Diagrams of NF-L structure, landmarks and epitopes. **A.** Diagram of the domain organization and epitope map of the human NF-L protein. **B.** Amino acid sequences including the Uman epitopes and the various peptides used. Core of UD2 epitope highlighted in red, region essential for UD1 binding highlighted in blue. The a and d positions of the hydrophobic heptad repeat are identified below the sequences. The region of the “stutter” where the normal heptad repeat breaks down is as indicated in the 333-344 region, with the addition of h, i, j and k amino acids. The charged i, i+4 residues are indicated with appropriate + and - signs above the sequence and instance is numbered in red.

In 2002 Norgren et al.^20^ described a panel of six mouse monoclonal antibodies made against purified mammalian spinal cord NF-L. Two of these antibodies, named 2.1 and 47.3, became the focus of interest as they sensitively detect NF-L in an ELISA.^21,22^ These two antibodies became key reagents in a commercial 96 well NF-L assay produced by Uman Diagnostics (Umeå, Sweden), the NF-Light^™^ ELISA. Uman Diagnostics was recently purchased by Quanterix (Billerica, MA) which markets this assay and a variant using the same two antibodies on the significantly more sensitive and convenient Single Molecule Array (Simoa^™^) bead-based platform. The Uman <u>47.3</u> antibody serves as the NF-L capture reagent and <u>2.1</u> as the detection reagent, which in the Uman kit are referred to as <u>UD1</u> and <u>UD2</u> respectively. Most of the recent interest in NF-L as a biomarker of CNS damage and disease is a result of the efficiency of these two antibodies in the Uman and related assays, especially as applied to human serum samples.

In the original study Norgren et al.^20^ showed that UD1 and UD2 both bound a chymotryptic fragment of NF-L identified as the entire ~320 amino acid α-helical rod region (**Fig. 1A**). Lower molecular weight chymotryptic fragments were bound identically by the two antibodies, suggesting that the relevant epitopes were close together. However these fragments was not characterized so the location of the epitopes was unknown. Competition studies showed that neither antibody inhibited the binding of the other to NF-L, suggesting that the antibodies bind to distinct and non-overlapping epitopes. Attempts at epitope mapping with nested peptides and by phage display were unsuccessful.^20^ Fully understanding a biomarker assay requires knowledge of exactly what it is detecting, focusing attention on the epitopes detected by the capture and detection antibodies. Such understanding may also allow the generation of improved second-generation NF-L assays, perhaps with novel or improved properties. Here we reveal the exact location of the epitopes for both Uman antibodies, and show that surprisingly they do not stain neurofilaments in healthy cells but strongly recognize degenerating and degenerated material. These findings stimulated us to generate a panel of novel NF-L antibodies to the Uman epitopic region which share these interesting neurodegeneration specific staining properties and which are briefly described here for the first time.

## Materials and methods

### Recombinant Constructs

We generated full length human NF-L, human NF-L rod region and human NF-L Coil 2 region, based on the sequence in <u>NP_006149.2</u>. All three constructs were made from DNA sequences from GenScript (Piscataway, NJ) which were codon optimized for expression in *E. coli*. The full length and the Coil 2 region constructs correspond to human amino acids 1-543 and 256-400 and are commercially available from EnCor, products <u>PROT-r-NF-L</u> and <u>PROT-r-NF-L-rct</u> respectively. The DNA sequences were inserted into the pET30a (+) vector between the *EcoR1* and *Sal1* cloning sites which adds N-terminal His-tag, S-tag and some other sequence, a total of 52 extra amino acids. A stop codon was inserted in the DNA sequence 5’ to the *Sal1* site. Constructs were expressed in *E. coli*. and purified in 6M urea using the Nickel affinity of the N-terminal His-tag by standard methods.

### Protein chemistry

Cleavage at tryptophan was performed with 3-Bromo-3-methyl-2-((2-nitrophenyl)thio)-3H-indole (BNPS-Skatole) and cleavage at cysteine residues was done using 2-nitro-5-thiocyanobenzoate (NTCB) as described.^23,24^ Both chemicals were obtained from Sigma-Aldrich (St. Louis, MO).

### Antibodies

**Supplementary table 1** lists relevant details of all antibodies used in this article, including the novel reagents presented here. For certain experiments 1mg amounts of purified <u>MCA-DA2</u>, <u>MCA-1B11</u> and <u>MCA-6H63</u> were directly labeled following the manufacturer’s instructions with the fluorescent dyes Alexa Fluor^®^ 488 or Alexa Fluor^®^ 647 using Invitrogen/Life Technologies Protein Labeling Kits and following the manufacturers protocol (cat# A10235, A20173).

### Peptides

We obtained thirtysix 20 amino acid peptides overlapping by 5 amino acids in 0.5mg amounts in 96 well format microtubes based on the entire human NF-L sequence in <u>NP_006149.2</u> (Sigma-Aldrich). Peptides were diluted in 50% acetonitrile/50% PBS to a concentration of 2.5mg/mL and 2mL of each peptide was arrayed in a 96 well plate. Then 100mL of the antibody to be tested at 1mg/mL was added in PBS plus 0.1% nonfat milk. The final concentration of peptide was ~23μM while the antibody concentration was about 6.6nM, so the peptide was presented at about 3,500 fold molar excess per antibody molecule. After 10 minutes the mix was transferred to an Immulon 4HBX plate previously coated overnight with 1μg/mL recombinant full length human NF-L in 50mM sodium bicarbonate pH=9.5 and then blocked with 5% nonfat milk in Tris buffered saline plus 0.1% Tween 20 (TBSt). After 1 hour at room temperature with shaking the plate was washed three times in TBSt, incubated for one hour with 1μg/mL goat anti-mouse alkaline phosphatase (Millipore Sigma Cat #AP124A), washed again three times in TBSt and signal was developed with p-nitrophenyl phosphate (pNPP Sigma Cat #N2765), the color reaction being quantified at 405nm.

Following the approximate localization of the Uman epitopes, Peptide 2.0 (Chantilly, VA) synthesized a 55 amino acids human NF-L peptide, amino acids 316-370 with the addition of a C-terminal cysteine to facilitate coupling to carrier proteins. After getting positive binding to this, Peptide 2.0 then synthesized NF-L 305-345, 316-357, 316-345, 328-357 and 341-370, each with an additional C-terminal cysteine (**Fig. 1B**). For direct binding experiments these were dissolved in TBSt diluted to 5mM and 100μL amounts were adsorbed onto Immulon 4HBX 50mM sodium bicarbonate pH=9.5 overnight at 4°C, then blocked in 5% nonfat milk in TBSt. Antibody incubations and signal development was as described above.

For peptide competition experiments we coated plates with 100μL of full length recombinant human NF-L at 0.5μg/mL in 50mM sodium bicarbonate pH=9.5 and then blocked with 5% nonfat milk in Tris buffered saline plus 0.1% Tween 20 (TBSt). The various peptides were each diluted to 5mM in TBSt and 100μL of each was added to the first well of a row of 10 wells on an ELISA plate, the remaining 9 wells filled with 50μL of TBSt. 50μL of the solution in the first well was serially diluted down the next 8 wells, the tenth row containing no peptide. 50mL of antibody diluted to 1.0μg/mL were added to each of the 10 wells and the plate was incubated for 1 hour at room temperature. Washing, secondary antibody incubations and signal development was as described above. This experiment tests binding of 0.5nM antibody to human NF-L in the presence of from 25,000nM to 0nM peptide.

Finally, GenScript synthesized thirty-five 25 amino acid peptides covering human NF-L 309-367. These peptides were obtained in 4mg quantities and were each dissolved in 1mL 50% distilled water/50% acetonitrile. Immulon 4HBX ELISA plates were coated overnight with 100mL per well of 0.5μg/mL full length recombinant human NF-L in 50mM carbonate buffer at pH=9.5. The next day the plate was blocked in 5% nonfat milk in TBSt for one hour at room temperature. The plate was emptied and 1μL of each peptide was added sequentially to three wells. 100μL of the relevant antibody was added at 1μg/mL and incubated for 1 hour at room temperature with shaking. Washing, secondary antibody incubations and signal development was as described above.

### Spinal Cord Injury

A total of ten adult female Sprague-Dawley rats (12-20 weeks of age) were obtained from Envigo (Hsd:Sprague Dawley^®^ SD, Indianapolis, IN). All procedures were approved by the Institutional Animal Care and Use Committee at the University of Florida (Protocol # 201807438) and are in accordance with National Institutes of Health Guidelines. Animals were housed individually in cages under a 12 hour light/dark cycle with *ad libitum* food and water. Animals were anesthetized with ketamine (90mg/kg) and xylazine (10mg/kg) via intraperitoneal injection. Before surgical procedure, sufficient depth of anesthesia was confirmed using a toe pinch resulting in lack of a change in heart rate, whisker twitch, and hindlimb withdrawal. Anesthesia was also continuously monitored during surgery and the animal was re-dosed with a one-third dose of the initial ketamine/xylazine dose when needed. Body temperature was maintained with a water recirculating heating pad. A dorsal incision was made from the base of the skull to the C6 region of the spine. Dorsal paravertebral muscles between C1 and C6 were incised and retracted. The posterior portion of cervical vertebrae was exposed, and a laminectomy was performed at the C4 level, preserving the facet joints and leaving the dura intact. The procedures for C4 midline contusion have previously been described.^25,26^ Briefly, the rat was suspended by clamps on the C3 and C5 vertebrae. Rats were subjected to a single contusion injury at the midline, using a 2.5mm diameter impactor tip. A force of 200kdyn with 0 second dwell time was delivered using the Infinite Horizon Impactor (Precision Systems and Instrumentation, Lexington, KY). For dorsal hemilesion, the dorsal aspect of the C4 spinal cord was sectioned using a spring-scissor (Cat # 15006-09, Fine Science Tools). To ensure correct depth of the lesion and consistency across animals, a 1mm mark was made on the knife prior to lesioning. This process was repeated three times to ensure completeness of dorsal lesion. Subsequently the overlying muscles were sutured with sterile 4-0 Webcryl and the overlaying skin was closed using 9mm wound clips. Animals were maintained on a heating pad until alert and awake. Animals were monitored on a daily basis for signs of distress, dehydration, and weight loss, with appropriate veterinary care given as needed. Animals received buprenorphine (0.03mg/kg, b.i.d.), meloxicam (1mg/kg, q.d.), Baytril (5mg/kg, q.d.), and lactated Ringer’s solution (10mL/day, q.d.) for 48 hours post-injury.

### Cell and tissue staining

Rats were anesthetized with isofluorane, perfused with fresh 4% formaldehyde in PBS and tissues dissected out and stored overnight in fresh 4% formaldehyde at 4°C. They were then transferred to PBS containing 15% sucrose in PBS for ~2 days, then 30% sucrose in PBS for ~2 days. Frozen sections were cut at 15μM on a Leitz cryotome and stained using standard procedures. Specimens were imaged on a Zeiss Axioskop 2 fitted with a Diagnostic Instruments RTke digital camera, a Keyence BZ-X810 microscope or an Olympus FV3000 confocal microscope.

### Protease Experiments

Dried formaldehyde fixed 50mM floating sections of uninjured rat spinal cord were rehydrated in PBS and then treated with 0.25% trypsin EDTA solution (Gibco cat# 25-200-056) or 20mg/mL proteinase K (recombinant PCR grade, ThermoFisher Cat# EO0492) for 10, 20, 60, 90 and 120 minutes. They were then washed in PBS to remove enzyme and fixed in 4% formaldehyde for 5 minutes, followed by regular immunostaining. Control sections were treated with the way the only difference being the omission of the enzyme. Enzyme treated sections were imaged on a confocal microscope and the exact settings optimized for these specimens were then used to image the relevant controls.

### Antibody Generation

All procedures were performed according to the NIH Guide for the Care and Use of Experimental Animals. For monoclonal antibody production 12 week old BALB/c mice were obtained from Jackson Labs (Bar Harbor, ME) and were injected subcutaneously on the back with 100μL of a purified recombinant construct containing human NF-L amino acids 311-362, which includes the two Uman epitopes, at 1mg/mL mixed 1:1 with Freund’s complete adjuvant. Three weeks later mice were boosted with the same immunogen in Freund’s incomplete adjuvant and 10 days after that blood samples were screened by western blotting on rat spinal cord homogenates. Two mice with good reactivity to NF-L were boosted with the same immunogen. After two weeks they were given a final intraperitoneal boost and three days later they were euthanized and spleens aseptically removed. Spleen cells were fused to PAI-O myeloma cells^27^ by standard methods and fused cells were seeded into six 96 well culture dishes. 10 days later hybridoma supernatants were screened by ELISA and western blotting on recombinant human NF-L, and promising clones were tested on sections of contused spinal cord. Useful clones were subcloned and characterized in detail.

## Results

### Epitope Mapping of Existing Antibodies

We performed preliminary competition experiments with a set of 20 amino acid peptides based on the full-length human sequence of NF-L, in which each peptide overlapped the next by 5 amino acids. We tested the inhibition of binding of Uman antibodies UD1 and UD2 and various other available NF-L monoclonal antibodies to pure recombinant full length human NFL (NF-L 1-543) in an ELISA format. UD2 binding was strongly inhibited by peptide 22, amino acids 316-335 of the human NF-L sequence, which is just N-terminal to the second “stutter” in Coil 2 (**Fig. 1A, Supplement fig. 1A**). The previous peptide, peptide 2, showed weak inhibition suggesting that the overlapping 5 amino acids TLEIE are an important part of the UD2 epitope, which is therefore dependent on the peptide TLEIEACRGMNEALE, amino acids 316-330. The widely used antibody MCA-DA2 was strongly inhibited by peptide 30, amino acids 436-455 of the human sequence and to a lesser degree by peptide 31, amino acids 451-470 in the “tail” region (**Supplement fig. 1B**). The epitope for this antibody is therefore dependent on the sequence SYYTSHVQEEQIEVE, amino acids 441-455. No convincing peptide inhibition of antibody binding to full length human NF-L was seen with UD1 or MCA-1B11. A summary of these results is shown in **Supplement fig. 1C.**

We tested both Uman antibodies on full length recombinant human NF-L (1-543), on a rod construct (human NF-L 80-400) and on a Coil 2 construct (human NF-L 256-400, **Fig. 1A, 2A**). Both antibodies showed strong staining for all three constructs, also showing that potential post translational modifications of NF-L do not significantly affect binding of either antibody. The NF-L 256-400 construct contains a single tryptophan and a single cysteine residue (**Fig. 1A**). Cleavage of the construct C-terminal to tryptophan 279 with BNPS-skatole is expected to produce two fragments, an N-terminal 8.5kDa and a C-terminal 13.9kDa (**Fig. 2B)**. The identity of the 13.9kDa band as the C-terminal fragment was confirmed using the <u>α-IFA</u> antibody which binds to a highly conserved peptide found at the C-terminus of the rod region of all 10nm filament subunits, amino acids 380-400 in human NF-L.^28^ The identity of the N-terminal fragment was confirmed using <u>Ab18184</u> antibody reactive with the vector derived His-tag sequence. Both Uman antibodies and MCA-1B11 bound the 8.5kDa C-terminal fragment, firmly mapping all three antibodies to within NF-L 280-400. NTCB cleavage of NF-L 256-400 N-terminal to cysteine 322 is expected to produce an N-terminal 13.2kDa and a C-terminal 9.3kDa fragment. The identity of these two fragments were confirmed using α-IFA and an antibody to the vector derived N-terminal S-tag sequence, <u>MCA-1B63</u> (**Fig. 2C**). UD1 and MCA-1B11 stained the C-terminal 9.3kDa fragment. Despite several attempts UD2 did not stain either fragment, although it recognized the uncleaved protein. This suggests that the UD2 epitope is destroyed by cleavage at cysteine 322, consistent with our finding that the 316–335 peptide including the single NF-L cysteine residue inhibited binding of UD2 to NF-L.

**Figure 2:**
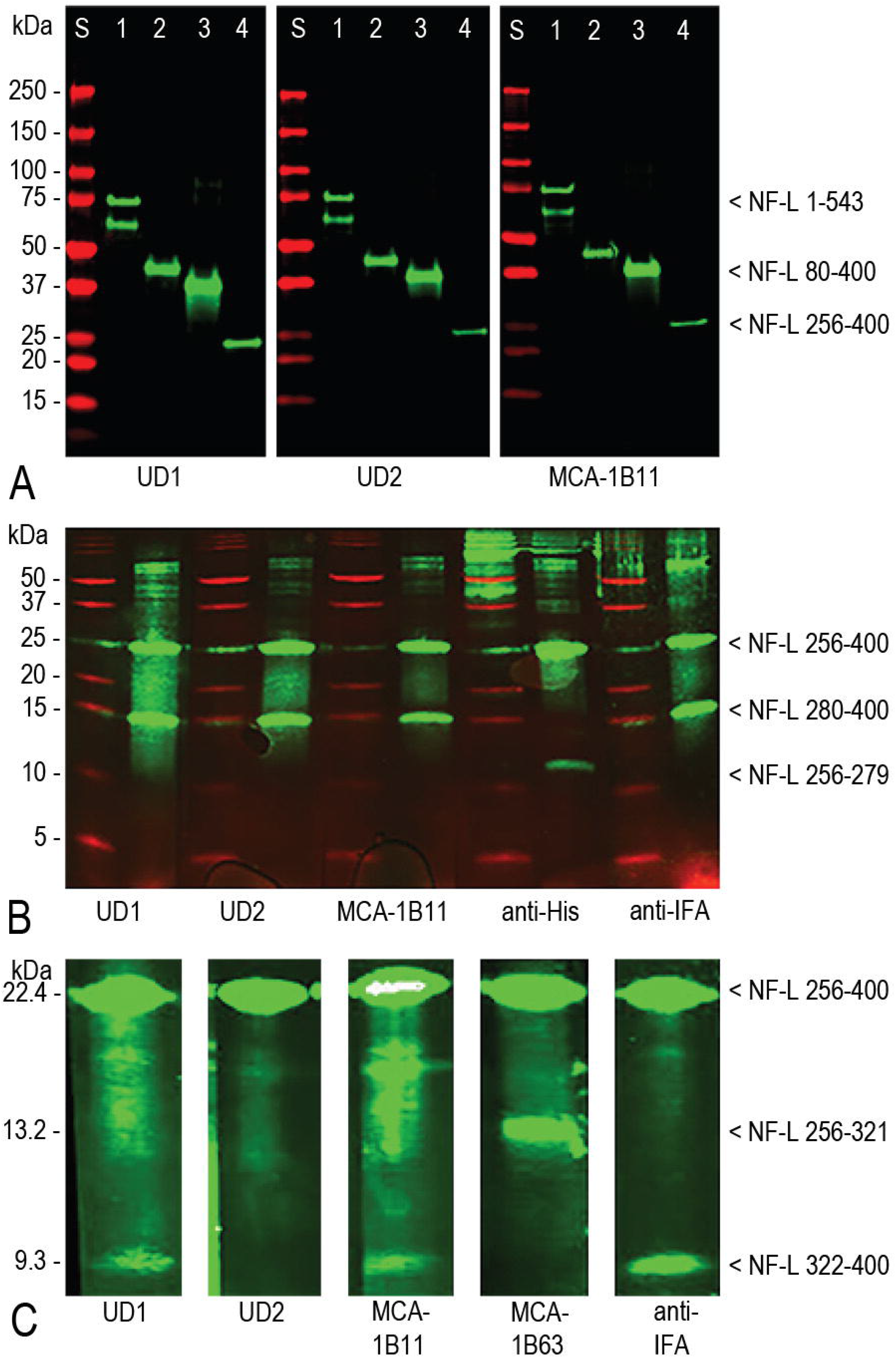
Western blotting studies. **A.** Western blotting of Uman and MCA-1B11. Lane S are molecular weight standards of indicated molecular weight. 1 is full length recombinant human NF-L, 1-543. 2 is a rod construct, NF-L 80-400, 3 is a proprietary immunogen used to generate novel antibodies, 4 is recombinant human NF-L Coil 2, amino acids 256-400. **B.** Partial protein cleavage of the 256-400 construct at tryptophan 279 produces two fragments of expected size 8.5kDa and 13.9kDa. All three antibodies tested bind to the C-terminal 13.9kDa band. **C.** Partial cleavage of NF-L 256-400 at cysteine 322, expected to produce an N-terminal 13.2kDa fragment and a C-terminal 9.3kDa fragment. Both UD1 and MCA-1B11 bind the C-terminal fragment while the epitope for UD2 is apparently destroyed. All three antibodies bind the uncleaved form of the protein.

The data suggested that the UD2 epitope was associated with cysteine 322, while UD1 and MCA-1B11 were C-terminal and likely close to this. We therefore synthesized peptide 316-370 of the human NF-L sequence with a C-terminal cysteine added (**Fig. 1B**). All three antibodies strongly bound this peptide either free, or coupled to MBS-ovalbumin on ELISA plates, or to the ovalbumin-peptide conjugate on western blots, localizing the epitopes for all three antibodies to within NF-L 316-370 (not shown). We then synthesized several smaller overlapping peptides, specifically NF-L 305-345, 316-357, 316-345, 328-357 and 341-370 (**Fig. 1B).** UD1 bound directly to 316-370 immobilized to ELISA plates and showed reduced binding to 316-357 and no binding to the other peptides **(Supplement fig. 2A)**. UD2 strongly bound to NF-L 316-370 and to a lesser degree to the three other peptides containing the 316-330 peptide (**Supplement fig. 2B)**.

We also tested the ability of each of the 5 peptides to inhibit antibody binding to recombinant human NF-L immobilized in an ELISA plate. The 316-370 peptide proved to be the most potent inhibitor of binding of both Uman reagents, suggesting that both are dependent on conformations not fully recapitulated by the shorter sequences. In line with the direct peptide binding results for UD1, 316-357 showed reduced inhibition while the other peptides were ineffective (**Supplement fig. 2C**). Also consistent with previous data UD2 binding was inhibited strongly by NF-L 316-370 and somewhat less efficiently by other peptides containing the 316-330 while other peptides were ineffective (**Supplement fig. 2D and insert**).

To further delineate the epitopes for these antibodies we generated a set of thirty-five 25 amino acid peptides staggered by one amino acid and spanning the sequence from 309 to 367. The results for UD2 showed that peptides which contain TLEIEACRGMNEALE, amino acids 316-330, all strongly inhibited antibody binding to NF-L as expected (not shown). However, inhibition disappeared with the loss of glutamic acid residue 318, allowing us to refine the core of the UD2 epitope to EIEACRGMNEALE, amino acids 318-330. Despite several trials under different conditions, we could not get convincing peptide inhibition of binding of UD1 to recombinant NF-L or direct binding of this antibody to any of the 25 amino acid peptides. We conclude that since UD1 bound NF-L cleaved at cysteine 322 on western blots (**Fig. 2C**), that the NF-L binding for this antibody is heavily dependent on sequence within 322-357 but that key components for strong binding extend beyond this region. The NF-L binding of UD2 is heavily dependent on NF-L 316-330 although, again, flanking sequences contribute to a more efficient interaction.

### Immunostaining of control cells and tissues

We stained 7-10 day old mixed neural cultures derived from E20 rat cortices with both Uman antibodies and compared the staining to that obtained with an affinity purified rabbit polyclonal antibody raised against rat NF-L amino acids 515-543, the C-terminal peptide of the NF-L tail. This antibody, <u>RPCA-NF-L-ct</u>, reliably produces strong and specific staining of neurons and their processes on rodent and other mammalian tissues. To our surprise both Uman antibodies failed to stain the majority of the clearly fibrillar neuronal profiles which were robustly stained with RPCA-NF-L-ct (**Fig. 3A,** UD1; **Fig. 3B,** UD2). Under high magnification we observed that staining with both Uman antibodies was primarily punctate and most of the material reactive with either Uman antibody was negative for RPCA-NF-L-ct. A few linear profiles were stained with the Uman reagents which in the majority of cases showed no staining for RPCA-NF-L-ct (**Fig. 3B**). Punctate and globular Uman positive profiles frequently appeared in discontinuous linear arrays which suggested that they may have originated from degenerated processes, in stark contrast to the well defined RPCA-NF-L-ct positive profiles (**Fig. 3C**). We then immunostained normal adult rat spinal cord and brain tissue sections with each of the two Uman antibodies and co-stained with RPCA-NF-L-ct. Both Uman reagents failed to stain the vast majority of cell bodies and processes which stained robustly with RPCA-NF-L-ct. We identified a few profiles in normal adult rat spinal cord which were positive for either UD1 or UD2 against a massive background of RPCA-NF-L-ct positive and UD1 and UD2 negative material (**Fig. 3D**, UD2 shown). We also noted a few rare neuron cell bodies in brain which stained with one or other Uman reagent but not with RPCA-NF-L-ct (**Fig. 3E, F**).

**Figure 3:**
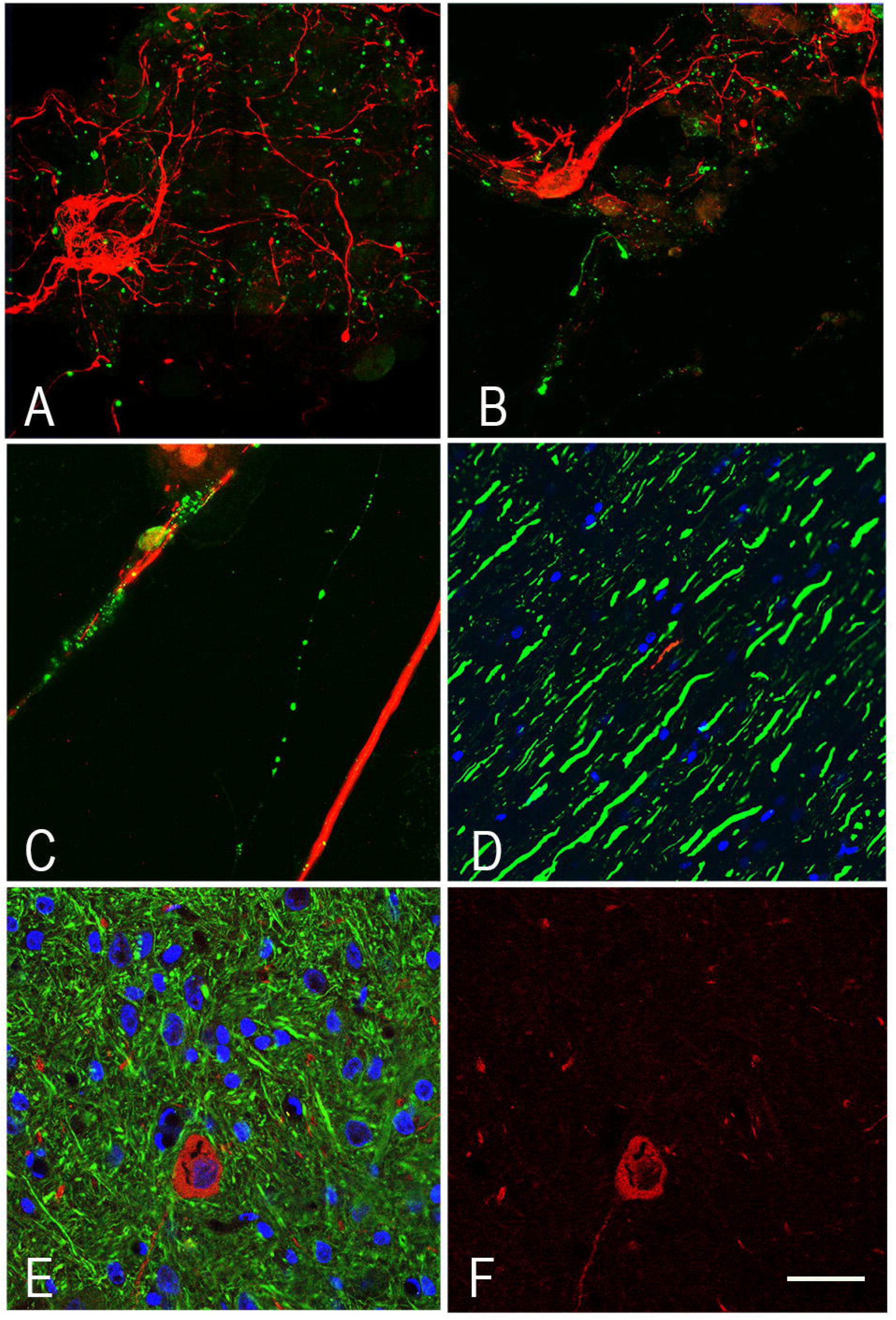
Immunofluorescence of control cells and tissues. **A**, **B**. 7-10 day neural cultures from E20 rats stained with UD1 (A) and UD2 (B) in green, both co-stained with RPCA-NF-L-ct in red. In both cases the majority of the Uman positive material is punctate and negative for RPCA-NF-L-ct while the RPCA-NF-L-ct antibody stains fibrillar profiles in cells with a typical neuronal morphology. **C** shows a region of a similar culture stained with UD2 in green and RPCA-NF-L-ct in red. The prominent linear array of Uman positive material is suggestive of the remains of a degenerated process, while the RPCA-NF-L-ct positive profile appears continuous and fibrillar. **D** shows a section of normal rat spinal cord stained with UD2 in red and RPCA-NF-L-ct in green. A single fiber negative for RPCA-NF-L-ct is revealed with the UD2 antibody. **E** and **F** show a region of brain stem from a control rat. One apparently unhealthy looking neuron and associated processes is positive for UD2 but not RPCA-NF-L-ct. Panel F shows the same specimen with only the Uman antibody. Blue in every case was staining with the DNA stain DAPI. Bar in figure F = 50μM for images A-C = 20μM for D = 40μM for E = 50μM

Given our convincing peptide binding and western blotting studies we were confident that both Uman antibodies were highly specific for NF-L, so we developed the hypothesis that both antibodies bound epitopes which were only accessible in degenerating and/or degenerated processes. There is in fact a significant amount of spontaneous neuronal death in neural cultures similar to those used here,^29,30^ and normal CNS tissues in adult animals reveal the occasional process or neuron which has undergone spontaneous degeneration.^31^

### Immunostaining of spinal cord tissue from rats with spinal cord injury (SCI)

We next evaluated spinal tissue from rats following sub-acute SCI, expected to produce large numbers of degenerating and degenerated neurons and their processes.^25,26^ Adult rats received a C4 midline contusion or C4 dorsal hemilesion, and tissues were harvested after 1, 3 or 5 days. [F5] A series of both longitudinal and coronal sections from the injured region and flanking tissue were evaluated along with similar sections from untreated control animals. Sections from the control animals showed robust RPCA-NF-L-ct staining with, as expected, virtually no staining with either Uman reagent (not shown). In marked and spectacular contrast coronal sections stained 1, 3 or 5 days after dorsal contusion or hemilesion show numerous profiles positive for the Uman antibodies which were mostly RPCA-NF-L-ct negative amongst a larger background of numerous RPCA-NF-L-ct positive, Uman negative fibers. At all three time points and in both contusion and hemilesion specimens apparently identical material was revealed with both Uman antibodies in the dorsal columns, dorsal corticospinal tracts and lateral funiculi both rostral and caudal to the lesion site (**Fig. 4A,** UD2 green, RPCA-NF-L-ct red). A few Uman positive profiles were seen in other regions of the cord including in the gray matter. Under higher magnification the profiles positive only for the Uman antibodies often appeared swollen, granular and diffuse, in contrast to the generally sharply defined axonal profiles positive for only RPCA-NF-L-ct (**Fig. 4B,** UD2, **Fig. 4C,** UD1). Longitudinal sections taken further from the lesion site revealed linear arrays of mostly swollen and discontinuous material negative for RPCA-NF-L-ct (**Fig. 5A,** UD1**, Fig. 5B,** UD2). The RPCA-NF-L-ct were compact and well defined in contrast to the generally more diffuse and less well defined Uman positive material (**Fig. 5A, 5B** inserts). These Uman positive aggregates were visible more than 1 cm from the lesion site in dorsal and lateral spinal cord tracts in animals given lesions 1, 3 and 5 days previously. Closer to the lesion site we saw a few examples of processes, usually swollen or sinusoidal, which stained for RPCA-NF-L-ct and one or other Uman antibody (**Fig. 5C**). Very close to the lesion site we frequently saw spherical and elongated profiles which were positive for both RPCA-NF-L-ct and one or other Uman antibody (**Fig. 6A**).

**Figure 4:**
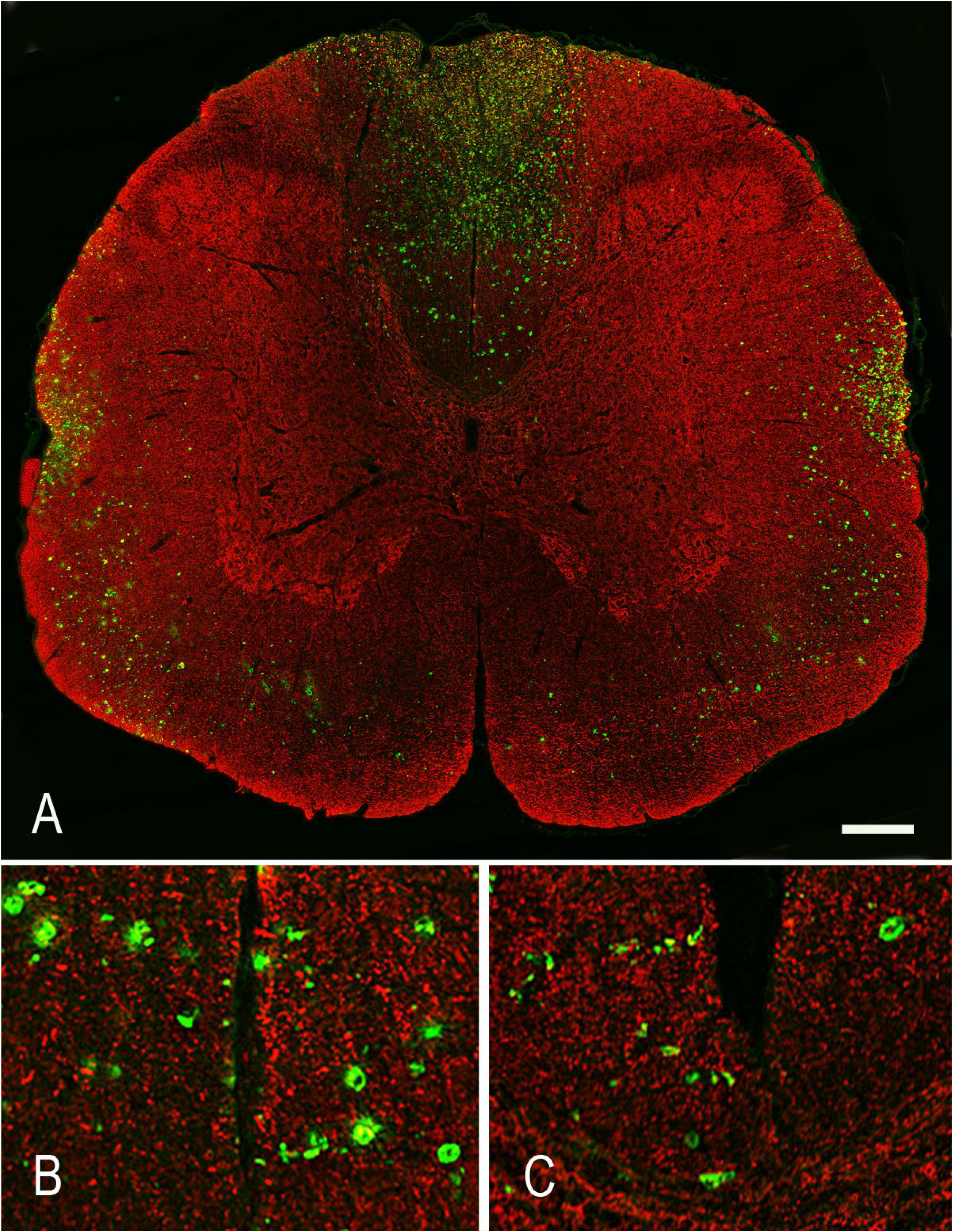
Immunofluorescence of coronal sections from contused 3 day rat spinal cord. **A** shows ancoronal section from a of rat given a contusion injury three days previously and stained for RPCA-NF-L-ct in red and UD2 in green. UD2 positive profiles are particularly obvious in the dorsal columns, corticospinal tracts and rubrospinal tracts, less abundant in the lateral and ventral funuculi and least abundant but not totally absent in the spinal cord gray matter. **B** shows a 6X magnified view of the region just above the central canal showing well defined RPCA-NF-L-ct axonal profiles in red in comparison to the more diffuse and generally non-overlapping UD2 profiles in green. **C** shows a 6X magnified view of the same region of a similar section stained with RPCA-NF-L-ct in red and UD1 in green. Bar in image A = 500μM, in B and C = 80μM.

**Figure 5:**
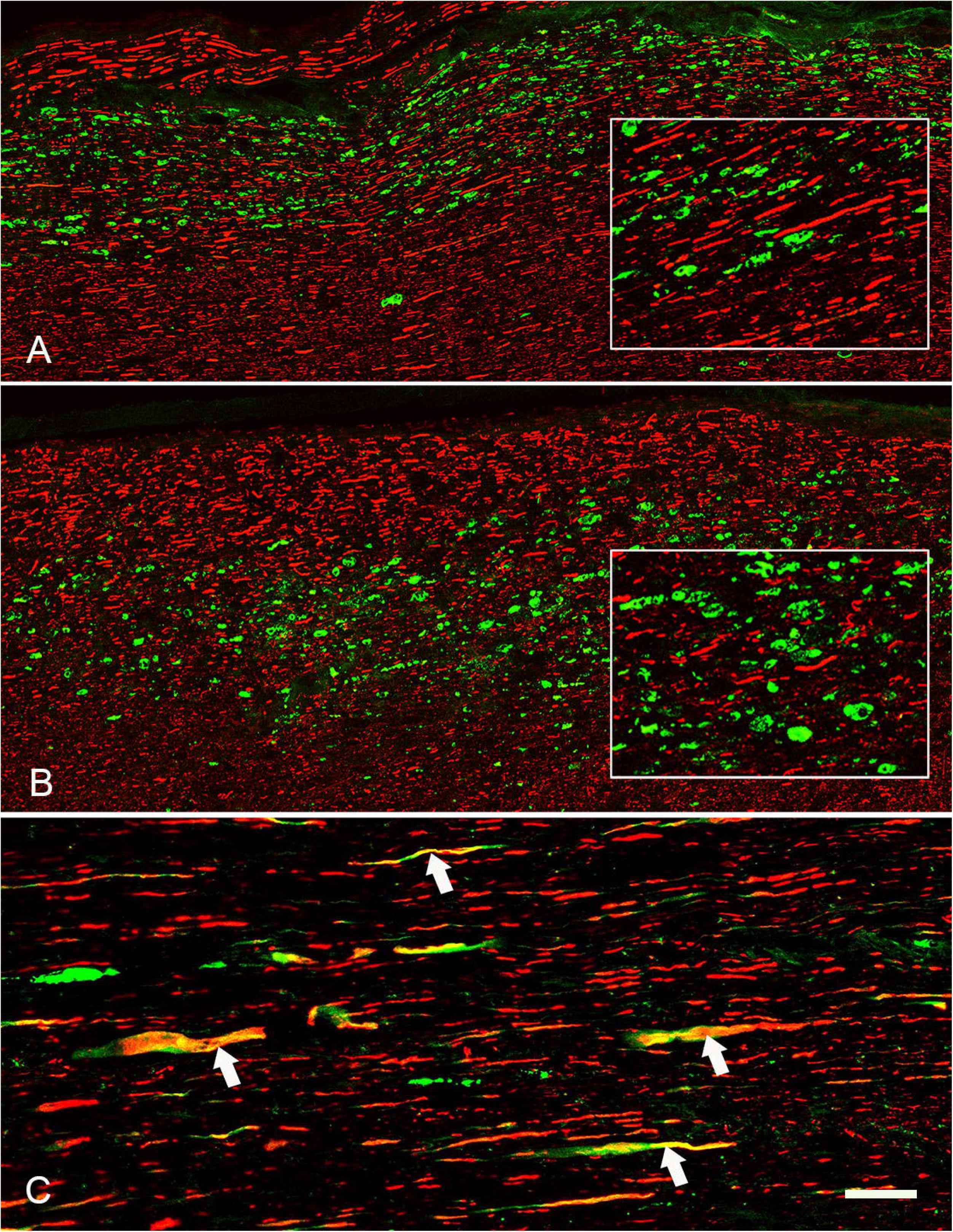
Immunofluorescence of longitudinal sections from 3 day contused rat spinal cord. Lateral rubrospinal and neighboring tracts some distance from the lesion. Linear arrays of profiles visualized with UD1 (**A**) and UD2 (**B**) are negative for RPCA-NF-L-ct. The RPCA-NF-L-ct profiles are generally well defined and continuous, the UD1 and UD2 profiles are mostly swollen, globular and discontinuous. Inserts show 3X magnified sections of each image. C shows a region close to a contusion lesion and shows swollen and apparently degenerating axonal profiles positive for RPCA-NF-L-ct in red but also show UD1 staining in green. Bar in figure C= 25μM, Bar = 50μM in A and B, 13mM in A and B inserts.

**Figure 6:**
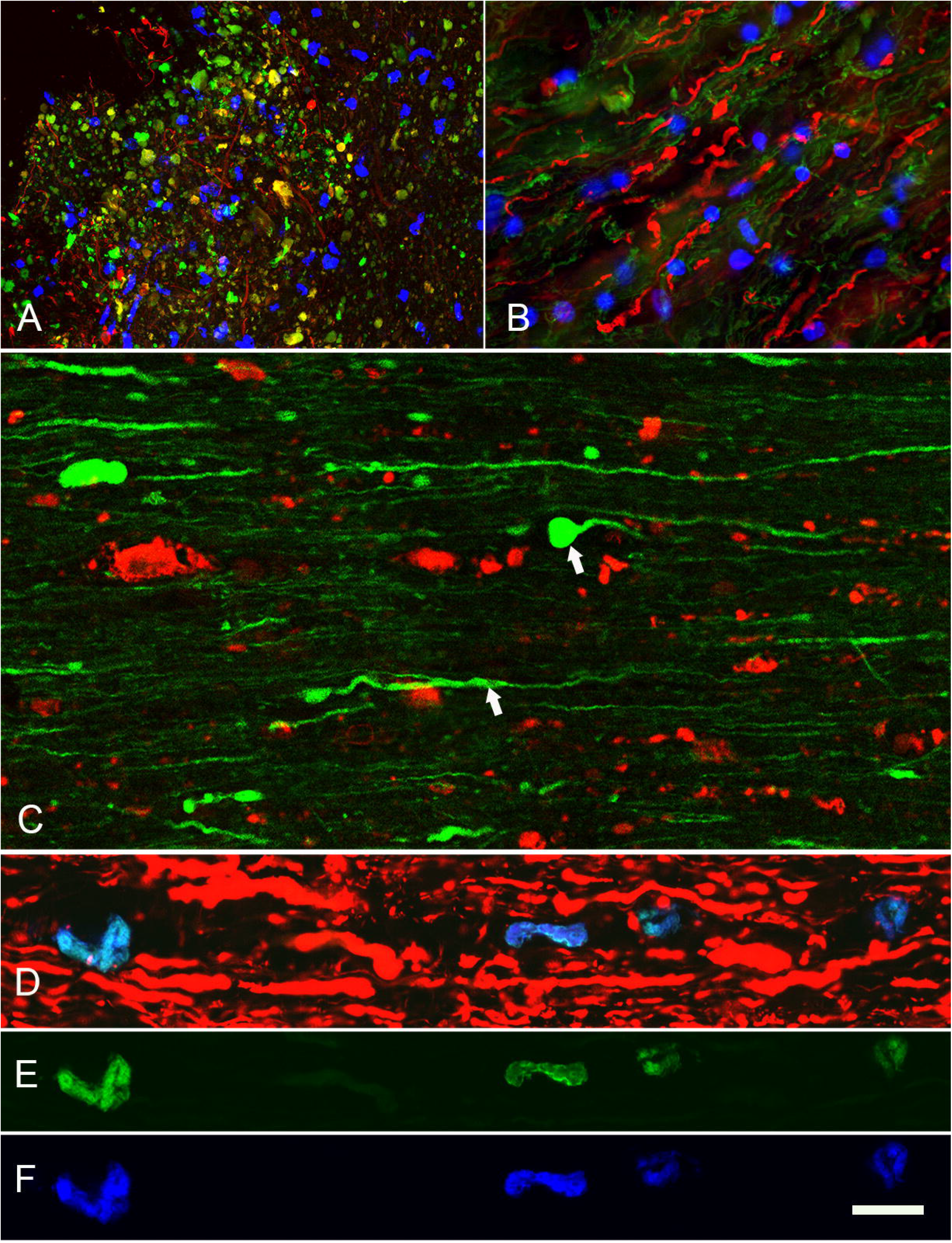
Further immunofluorescence of contused rat spinal cord. **A** shows a region at the lesion site of an animal given a contusion 3 days previously stained with RPCA-NF-L-ct in red and UD1 in green. Fibers positive for RPCA-NF-L-ct alone are visible amongst mostly globular profiles positive for both antibodies or only for UD1. **B** shows a region of adult mouse spinal cord which was allowed to sit at room temperature for 4 hours prior to fixation. Sections were then incubated with RPCA-NF-L-ct in green and MCA-1B11 in red. Note prominent beaded, discontinuous, sinusoidal and apparently helical MCA-1B11 positive profiles which are negative for RPCA-NF-L-ct, while RPCA-NF-L-ct staining is attenuated. **D-F,** a longitudinal section of spinal cord from an apparently healthy control rat was searched to find what we proposed are spontaneously degenerating or degenerated nerve fiber as in **Fig. 3D**. **D** shows typical axonal profiles stained with RPCA-NF-L-ct in red. A linear group of globular profiles which were negative for RPCA-NF-L-ct but positive for MCA-1B11 (**E,** green) directly coupled to Alexa Fluor 488 (green) were also identically positive for MCA-6H63 coupled to Alexa Fluor 647 (**F,** blue). MCA-1B11 and MCA-6H63 bind distinct hidden NF-L epitopes building confidence that the objects stained are indeed degenerating or degenerated axons. Bar in D = 10μM.

We directly labeled the NF-L antibody MCA-DA2 with Alexa Fluor-488 which allowed double label immunofluorescense with each of the Uman reagents and this mouse antibody. As shown above the MCA-DA2 epitope is within the C-terminal tail of NF-L but distinct from the region recognized by RPCA-NF-L-ct. Like RPCA-NF-L-ct, MCA-DA2 recognizes both normal and damaged neuronal processes, showing helical and typical retraction bulb endings close to the lesion site (**Fig. 6C**). Also like RPCA-NF-L-ct, MCA-DA2 does not stain most of the Uman positive material. We also obtained mouse monoclonal antibody 4F8 described by Rutherford et al.^32^ which recognizes a phosphorylation site in the NF-L tail region centered on serine 473, distinct from the MCA-DA2 and RPCA-NF-L-ct epitopes. This antibody also failed to bind the degenerated Uman positive material (not shown).

The obvious conclusion from these observations is that neuronal processes which were originally immunoreactive with certain NF-L tail antibodies lose this immunoreactivity while Uman epitopes concomitantly become unmasked. Some processes express both Uman and NF-L tail epitopes (**Fig. 5C**), but the relative paucity of these suggests that the transition from one state to the other is rather rapid. Obviously degenerated material then expresses only the Uman epitopes. The obvious hypothesis to explain these findings is that degeneration activates proteases which remove or destroy the NF-L tail epitopes and concomitantly expose the Uman epitopes.

### Protease Experiments

We incubated standard formalin fixed coronal sections of uninjured rat spinal cord with proteinase K or trypsin for various times. Control tissue sections which were treated with appropriate protease buffer lacking protease showed strong staining with RPCA-NF-L-ct and essentially no staining with either Uman antibody (**Figs. 7A and B, E and F**). In sharp contrast, trypsin or proteinase K treatment resulted in the generation of Uman antibody positivity with both antibodies, while staining for RPCA-NF-L-ct was diminished (**Figs. 7C and D, G and H**). We also dissected spinal cord tissues from an untreated mouse and left them at room temperature for 4 hours after sacrifice. The material left at room temperature contained numerous Uman positive linear objects including striking beaded and helical profiles, while the staining for RPCA-NF-L-ct was attenuated (**Fig. 6B**), while control tissues fixed immediately post mortem showed essentially no Uman positive material. Both experiments support our hypothesis that proteases expose the Uman epitopes concomitantly with the destruction or wash out of RPCA-NF-L-ct epitopes.

**Figure 7:**
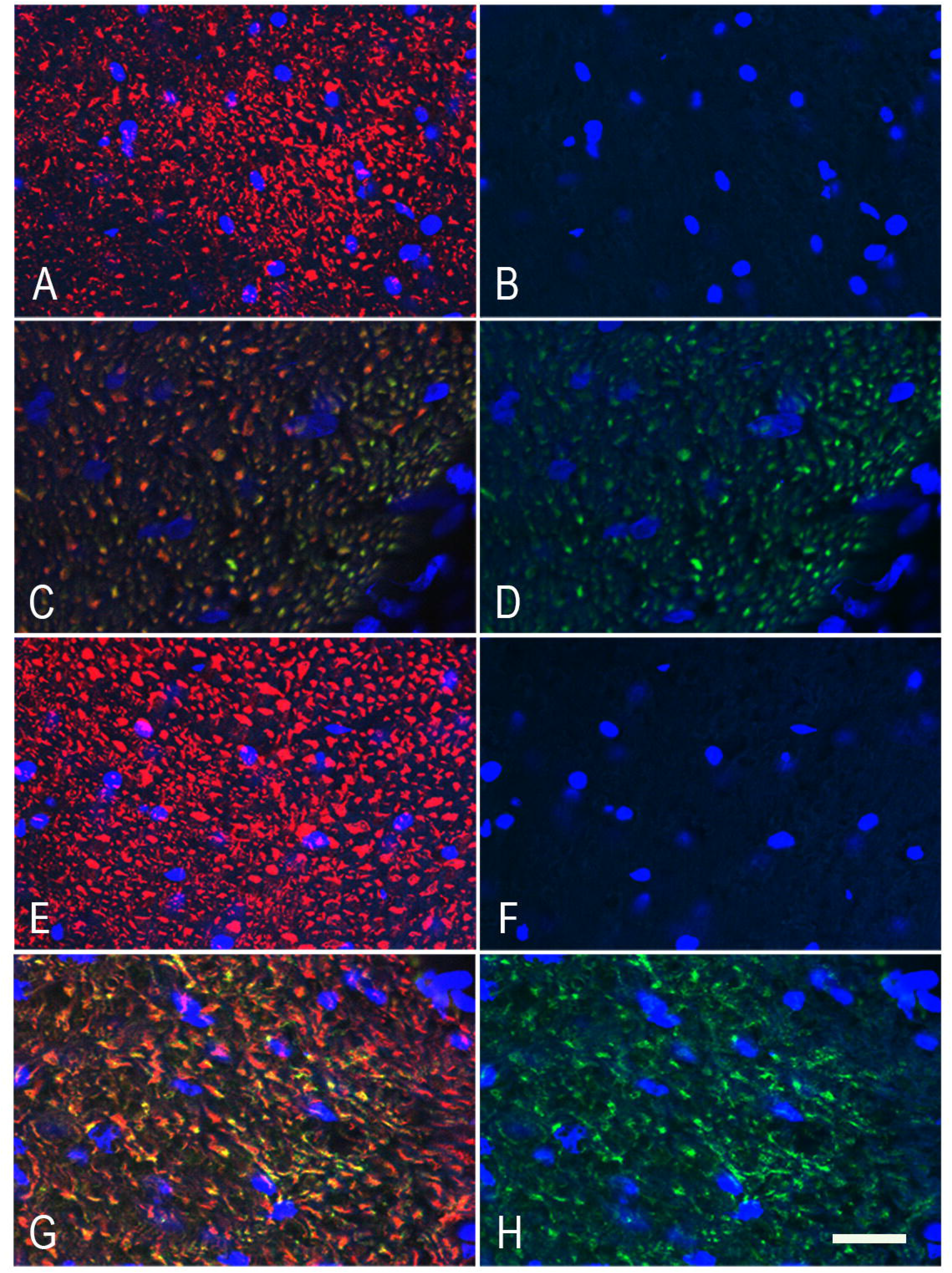
Protease experiments. Coronal sections of uninjured control rat spinal cords were treated with 0.25% trypsin or with buffer control lacking trypsin under exactly the same conditions. **A** shows a fiber tract after 10 minutes in control buffer then stained with RPCA-NF-L-ct in red and UD1 in green, while **B** shows just the UD1 signal. **C** and **D** shows a similar region incubated exactly as the section in **A** and **B** but with the addition of 0.25% trypsin. **C** and **D** were imaged on a confocal microscope and identical laser and software settings were then used to image **A** and **B**. Following enzyme treatment there is a weaker RPCA-NF-L-ct signal and a significant UD1 signal. **E** and **F** show similar control data for UD2 (green) and RPCA-NF-L-ct (red) on a similar section from an uninjured animal, and **G** and **H** show the staining of the same two antibodies following 10 minutes treatment in trypsin. Bar in H = 20μM.

### Generation and preliminary characterization of novel degeneration specific antibodies

We raised antibodies against a proprietary recombinant construct including both Uman epitopes and some flanking sequence, slightly different from the 316-370 peptide, at human NFL 311-362 (**Fig. 1B**). As expected, this immunogen was strongly reactive with both Uman and MCA-1B11, consistent with our conclusions concerning the location of the epitopes for these three reagents (**Fig. 2A**, third lanes). The novel monoclonal antibodies gave expected signals by ELISA, western blots of mammalian spinal cord extracts and on recombinant human NF-L constructs (not shown). Characterization of mouse monoclonal antibodies to 311-362 revealed clones which could be classified into at least four different groups, dubbed Classes I to IV. Class I includes UD1 and MCA-1B11 which were antibodies which could not be epitope mapped by competition using 25 amino acids peptides, bound to 316-370 immobilized in ELISA wells, and bound less well to 316-357. Class II antibodies included UD2 and several novel mouse monoclonal reagents including <u>MCA-1D44</u>. These showed strong binding to NF-L 316-370, 305-345, 316-357, 316-345 and more marginal or no binding for other peptides. These findings suggest that these antibodies bind epitopes similar to or identical to that of UD2. Class III antibodies such as MCA-6H63 bind to the N-terminal of the 311-362 immunogen but show no binding to other peptides including the 316-370 peptide. Their epitopes are therefore heavily dependent on NF-L 305-310, distinct from both UD1 and UD2 (**Fig. 1B**). Class IV antibodies such as antibody 6D112 bound to all peptides including the 341-357 sequence. We also generated affinity purified rabbit and chicken polyclonal antibodies to the 311-362 peptide which we named <u>RPCA-NF-L-Degen</u> and <u>CPCA-NF-L-Degen</u> respectively.

### Preliminary immunostaining results with novel antibodies

We focused on the Class I monoclonal antibody MCA-1B11, the Class II MCA-1D44 and Class III MCA-6H63 which all worked well on western blots and in peptide binding studies. Using these antibodies, we were able to replicate all the immunfluorescence findings we made with UD1 and UD2. All three novel reagents showed degeneration specific binding on the experimental SCI sections (see **Fig. 6B, 6E** for MCA-1B11, **Fig. 6F, 8E** for MCA-6H63, **Fig. 8F** for MCA-1D44).

**Figure 8:**
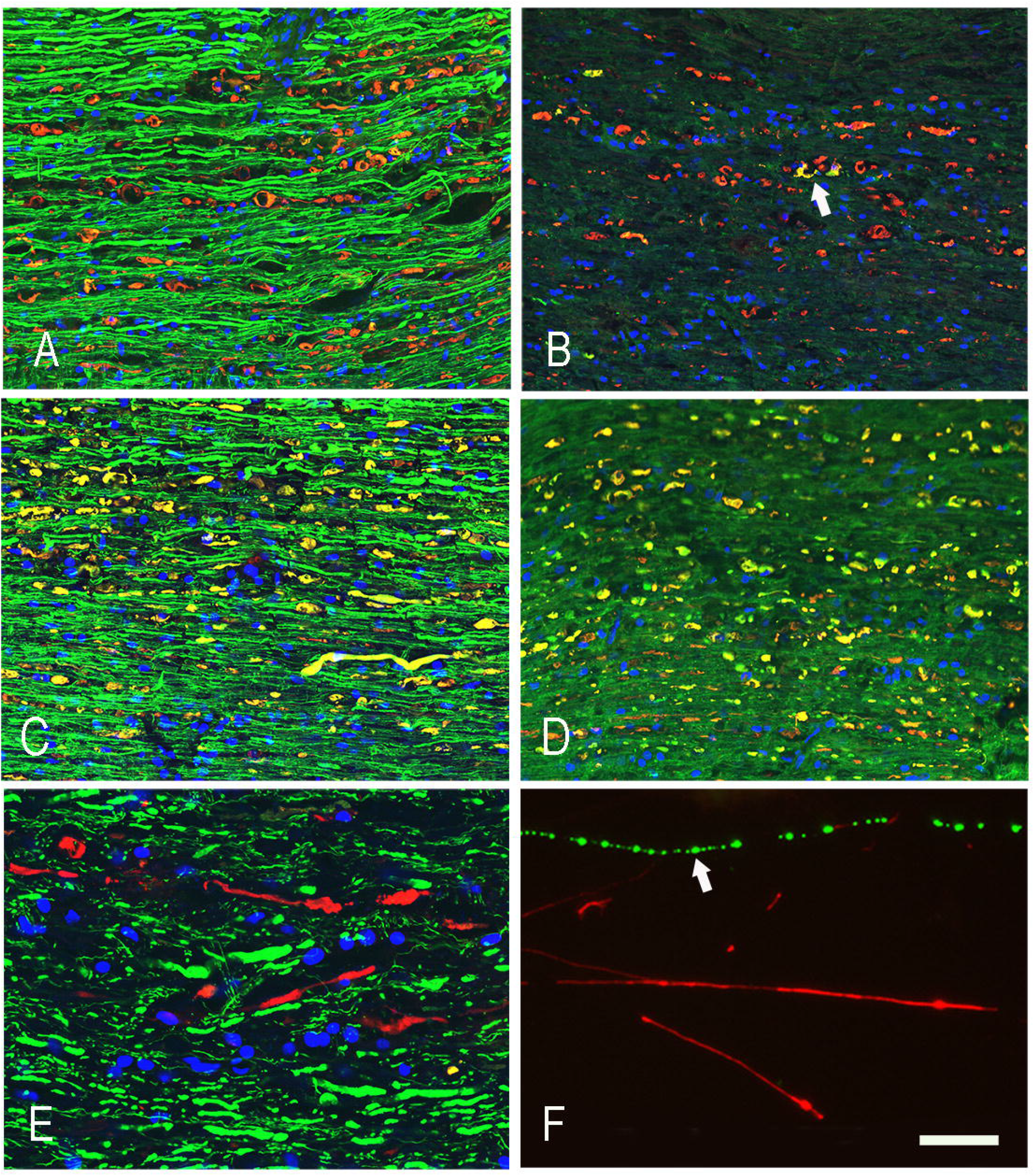
Immunocytochemical studies with novel antibodies. **A-E** Section of spinal cord from rat given a contusion 3 days previously and stained with RPCA-NF-L-Degen in red and, in green the following: **A** MCA-3H11 NF-M tail antibody. **B** is MAPt antibody MCA-5B10. **C** shows phospho-NF-H KSP tail sequences with MCA-NAP4, **D** shows MCA-2E3, mouse monoclonal to α-internexin. Clearly some epitopes are destroyed or washed out while others remain associated with the degenerated material. **E** shows longitudinal rat SCI tissue stained with MCA-6H63 in red and counterstained with RPCA-NF-L-ct in green. Despite having an epitope distinct from both UD1 and UD2 the MCA-6H63 antibody also clearly stains degenerating processes and not healthy processes. **F** shows MCA-1D44 staining of mixed neural cultures in green counterstained with RPCA-NF-L-ct in red. As with the Uman reagents (**Fig. 3C**), MCA-1D44 fails to stain what are apparently healthy neuronal processes but does stain linear sets of aggregates which must have originated from degenerated processes. Bar in panel H = 20μM, applicable to all images.

In our original studies we noted a few very rare Uman antibody positive fibers in spinal cord and brain sections of apparently healthy control animals (**Fig. 3D-F**). The availability of a panel of monoclonal antibodies with distinctly different epitopes allowed us to readdress this issue. We directly labeled the Class I monoclonal antibody MCA-1B11 with Alexa Fluor^™^ 488 and the Class III monoclonal antibody MCA-6H63 with Alexa Fluor^™^ 645. Control longitudinal sections of healthy spinal cord tissues revealed a few profiles which were negative for RPCA-NF-L-ct but were recognized strongly by both antibodies with an apparently identical staining pattern (**Fig. 6D-F**). This finding builds confidence that the few Uman antibody positive profiles in control tissues share at least two distinct hidden NF-L epitopes and so are very unlikely to be staining artefacts.

The novel rabbit and chicken antibodies raised against the 311-362 peptide showed, like the original Uman antibodies, showed strong staining of degenerating and degenerated material in SCI cords but little or no reactivity with control section from uninjured animals (**Fig. 8A-D** for RPCA-NF-L-Degen). Given the availability of these reagents we were able to identify degeneration induced aggregates which we could then costain with monoclonal antibodies providing a novel way of probing the axonal degradome. <u>MCA-3H11</u>, a monoclonal antibody directed against a sequence in the C-terminal tail of NF-M,^33^ showed strong axonal staining like RPCA-NF-L-ct in green but only weak staining of the inclusions identified with affinity purified rabbit polyclonal antibody to NF-L 311-362 (RPCA-NF-L-Degen) in red. (**Fig. 8A**) This suggests that the NF-M tail epitope, like the comparable region of NF-L, is either destroyed by proteolysis or washes away on degeneration. **Fig. 8B** shows staining for MAP-t with mouse monoclonal <u>MCA-5B10</u> which binds a peptide in the MAP-t core domain (green) counterstained with RPCA-NF-L-Degen in red. Surprisingly, some aggregates stain strongly for MAP-t while others do not. Possibly MAP-t washes out of the aggregates more quickly that the NF-L fragments. **Fig. 8C** shows a similar section stained with <u>MCA-NAP4</u> in green and RPCA-NF-L-Degen in red. MCA-NAP4 reacts with the phospho-KSP repeats of NF-H,^34^ and strongly reacts with the RPCA-NF-L-Degen positive inclusions. Apparently this region of NF-H does not readily wash out of the degenerated material. **Fig. 8D** shows staining for <u>MCA-2E3</u>, mouse monoclonal antibody to α-internexin, showing variable staining of the RPCA-NF-L-Degen positive inclusions. The epitope for this antibody lies within the C-terminal 166 amino acids of rat α-internexin equivalent to the C-terminal 160 amino acids of the human molecule,^35^ the entire tail region and the C-terminal part of Coil II. Future studies will follow up on these various findings.

Finally **Fig. 8E** shows longitudinal rat SCI tissue stained with Class III antibody MCA-6H63 in red and counterstained with RPCA-NF-L-ct in green. Despite having an epitope N-terminal to and distinct from those of UD1 and UD2 the MCA-6H63 antibody also clearly stains degenerating and not healthy processes. **Fig. 8F** shows Class II MCA-1D44 staining of mixed neural cultures in green counterstained with RPCA-NF-L-ct in red. As with the Uman reagents MCA-1D44 fails to stain what are apparently healthy neuronal profiles but does stain linear sets of aggregates which must have originated from degenerated processes.

## Discussion

The data presented contains one very surprising finding, that both Uman antibodies and our novel antibodies against the Uman epitopic region all fail to stain cells and tissues like other typical NF-L antibodies. With a few rare exceptions these antibodies did not bind NFs in healthy tissues which were strongly positive with other well characterized NF antibodies (**Figs 3–8**). Clearly, they are highly specific for NF-L as evidenced by their convincing binding to native mammalian and recombinant NF-L on western blots (**Fig. 2**), and by direct and competitive peptide binding experiments (**Supplement figs 1 and 2)**. We propose that during degeneration, proteolysis of NFs is induced and the Uman related epitopes become exposed and available for antibody binding. The concomitant loss of reactivity with RPCA-NF-L-ct, MCA-DA2 and 4F8 presumably means that certain C-terminal tail epitopes of NF-L are either degraded or removed. The Uman reagents and the novel antibodies described here therefore uniquely allow the identification of degenerated and degenerating neuronal profiles while RPCA-NF-L-ct, MCA-DA2, 4F8 allow the simultaneous visualization of healthy neuronal profiles. Neuronal profiles positive with both NF-L tail and Uman type antibodies have presumably been caught undergoing degeneration.

In line with our hypothesis, when we induced neurodegeneration by experimental SCI, we noticed a spectacular change in the staining pattern produced by both Uman antibodies with very large numbers of positive processes. Furthermore, most of these profiles were swollen, beaded, diffuse, sinusoidal or discontinuous in appearance, all hallmarks of degeneration. Our hypothesis is further strengthened by our finding that the Uman epitopes can be readily revealed by experimental protease treatment of tissue sections from healthy animals with no neural injury (**Fig. 7**), or by allowing tissues to naturally degrade (**Fig. 6B**).

The central region of 316-357 peptide contains the second Coil 2 “stutter”, a stretch of 11 amino acids which do not fit the typical hydrophobic heptad repeat pattern and the UD1 epitope includes this region (**Fig. 1B**). Structural data on the highly homologous region of the closely related protein vimentin reveals a continuous α-helical coiled coil, the stutter merely introducing a slight kink.^16^ The AlphaFold2 and RoseTTAFold structural modeling algorithms both predict that amino acids 328-357 of NF-L are part of a continuous elongated α-helix.^36,37^ The UD1 antibody cannot be inhibited from binding to NF-L by any 25 amino acid peptide tested and longer constructs such as 316-357 showed reduced inhibition when compared to the even longer 316-370 (**Supplementary fig. 2C**). This suggests that UD1 likely binds to an outside face of the slightly kinked region of the α-helical coiled coil structure, interacting with a series of amino acids which are not adjacent in the primary sequence. UD2 in contrast has a more linear epitope since amino acids 318-330 effectively but not fully inhibit UD2 binding to NF-L. (**Supplementary fig. 2D)**.

Three features of the NF-L region contains both Uman epitopes suggest that it may be functionally important. Firstly, experimental addition of three appropriate amino acids to “repair” the stutter 2 sequence in vimentin to form a typical heptad repeat produces a construct which still forms dimeric and multimeric assemblies.^38^ However it does not assemble into 10nm filaments suggesting that this region has an important role in higher order filament assembly. Secondly the Coil 2 second stutter is highly conserved in all 10nm filament subunits including the otherwise quite divergent keratins and nuclear lamins, suggesting some fundamental importance. Thirdly this region includes at NF-L 323-344 five tandemly arranged instances of amino acids alternating in charge separated by 3 amino acids (**Fig. 1B**). 10nm filament subunits are rich in instances where an amino acid at position i in an α-helical coiled coil sequence is followed by one of opposite charge at position i+4.^39^ However a stretch of five of these in tandem is very unusual and this pattern is conserved in all non-epithelial cytoplasmic 10nm filament subunits. The entire NF-L Coil 2 sequence contains 145 amino acids and 15 of the i, i+4 type sequences. Strikingly 5 are in the 21 amino acids of the NF-L 323-344 central to the Uman epitopic region. Letai and Fuchs^39^ suggested that such charge pairs may form stabilizing intramolecular interactions but may also be important in directing the formation of larger multimeric structures, perhaps by conformational changes from intra to inter-molecular charge interactions. None of these charged residues are in the a or d positions of the α-helical coiled coil sequence and so all face outwards from the α-helical dimer, potentially accessible for antibody binding and assembly interactions. Clearly filament assembly must be directed by specific protein-protein interfaces between NF-L dimers, tetramers or other smaller complexes. It is therefore not implausible that certain of these interfaces may be accessible to antibodies on denatured, partially assembled or proteolyzed filaments but inaccessible in fully assembled filaments. Our findings may also help explain why the Uman NF-Light^™^ assay and derivatives are so effective. The relevant region of NF-L is clearly proteolytically stable (**Figs. 6B, 7**), and the numerous charged residues likely result in great solubility.

The new antibodies described here and the original Uman reagents therefore have great utility beyond ELISA type assays. It will be interesting to apply these reagents to the numerous existing animal models associated with axonal loss such as amyotrophic lateral sclerosis, multiple sclerosis, traumatic CNS injury and many others. They are also useful reagents to identify spontaneous neuronal death in healthy animals. It seems likely that these antibodies would be useful for tracing fibers following surgical or chemical lesioning. Much evidence suggests that loss of axonal integrity is the major problem in a variety of CNS damage and disease states.^1–3^ Our reagents may become useful in further classifying the stages of axonal degeneration in combination with antibodies to for example amyloid precursor protein and other axonal injury markers.^3^ The NF-L epitopes we have elucidated are defined by amino acid sequences which are 100% identical in humans, apes, monkeys, rodents, pigs, dogs and apparently all mammals, so our new and the original Uman antibodies should be applicable to studies of any mammalian species or disease model. We must be cautious however, our immunocytochemical findings were made on mildly fixed material which may preserve the epitope masking we have discovered. Procedures such as epitope retrieval methods which work by partially denaturing proteins might expose the Uman type epitopes in assembled NF.

We note that Das et al.^40^ recently described two novel NF-L monoclonal antibodies, ADx206 and ADx209, for use in NF-L ELISA type assays. ADx209 was mapped to LQELEDKQNAD, amino acids 333-343, which corresponds to the stutter 2 region, making this a Class I antibody (**Fig. 1B**). ADx206 was mapped to VRAAKDEVSESRRL, amino acids 298-311, a few amino acids N-terminal to our Class III antibody MCA-6H63. It will be interesting to see if the ADx antibodies share the degeneration specific staining pattern described here. The Das et al. article also looked at various NF-L assay standards to which we can add our NF-L Coil 2 construct, amino acids 256-400, which is much smaller than those studied and perhaps more convenient. Finally, our newly generated reagents may obviously be used to develop NF-L ELISA type assays complementary to those made using existing reagents.

These findings raise many interesting questions. Is the Uman positive material we saw at 1 day post injury still there at 5 days, or has some or all of it been removed by 5 days? If the Uman positive material is rapidly removed this would mean that the Uman positive material seen at 5 days is the result of secondary neurodegeneration. In a previous study of experimental rat SCI we noted an acute peak of release of pNF-H into blood in the first 24 hours after injury followed by a much larger and more sustained release peaking at about 3 days which we speculated may have resulted from secondary degeneration^4^. Which proteases generate the Uman epitope containing NF-L fragment and where exactly do they cleave the NF-L molecule? Are different NF-L fragments generated in necrotic, apoptotic or other kinds of cell death? Does the Uman epitopic region get incorporated into any of the numerous kinds of inclusions associated with various neurodegenerative diseases? Does the hidden epitopic region have any function in activating inflammation or in appropriate circumstances regeneration? What is the half-life of the Uman positive material following degeneration and what mechanisms remove it? If our speculation is correct and the Uman epitopic region is a binding site important for higher order filament assembly, what does it bind to? This may be homotypic binding or alternately heterotypic, in which case the putative Uman region binding partner might also be hidden in assembled NF. Would a reagent based on the 311-362 peptide specifically prevent NF-L from forming NFs in vitro and in vivo? If so, these findings may also have some therapeutic implications for Charcot Marie Tooth disease forms caused by NF-L mutations. How does this method of visualizing neurodegeneration compare to other methods, such as TUNEL, Fluorojade and others? In conclusion, there are clearly numerous avenues for further investigation.

## Supporting information

Supplement Description

## Funding

Supported partially by grants 1R01HL139708-01A1 and 1 R01 HL153140-01 to David Fuller. EnCor Biotechnology Inc. also provided funds for this project from company income.

## Acknowledgements

Marda Jorgensen made many useful comments on earlier versions of the manuscript. We also thank Regina Shaw, Yong Wang and Vedrana Marin for the generation of immunogens and preliminary characterization of MCA-DA2, MCA-1B11 and other NF-L antibodies utilized here. We also thank Benoit Giasson, of the University of Florida, for NF-L antibody 4F8.

## Competing interests

Gerry Shaw is founder and majority owner of EnCor Biotechnology and may gain income and equity from the sale of reagents described in this article. Irina Madorsky, Ying Li and YongSheng Wang are full time employees of EnCor Biotechnology and may also benefit from indirect income growth. Sabhya Rana and David Fuller are full time employees of the University of Florida and have no competing financial conflicts of interest.

