## Supplementary figures and images for "Uman Type NF-L Antibodies Are Effective Reagents for the Imaging of Neurodegeneration"

### Supplement Figure 1

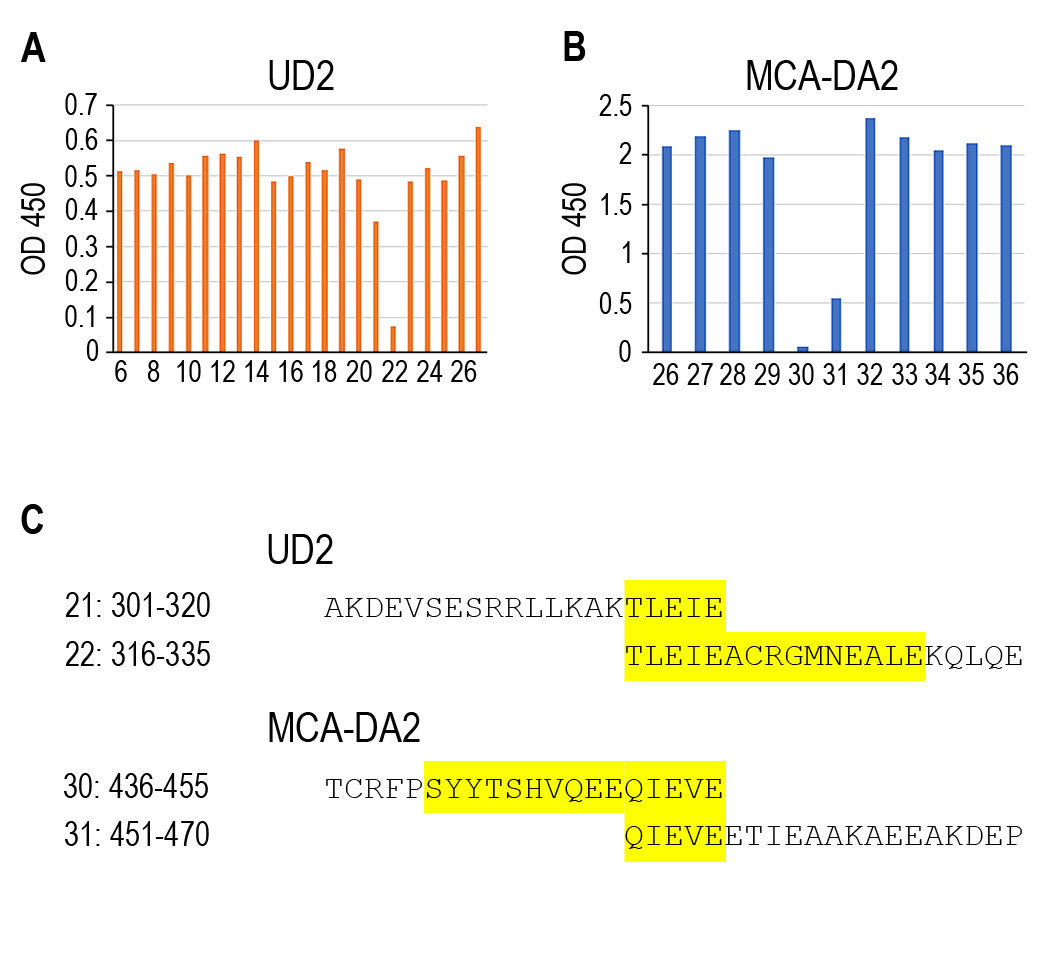

### Supplement Figure 2

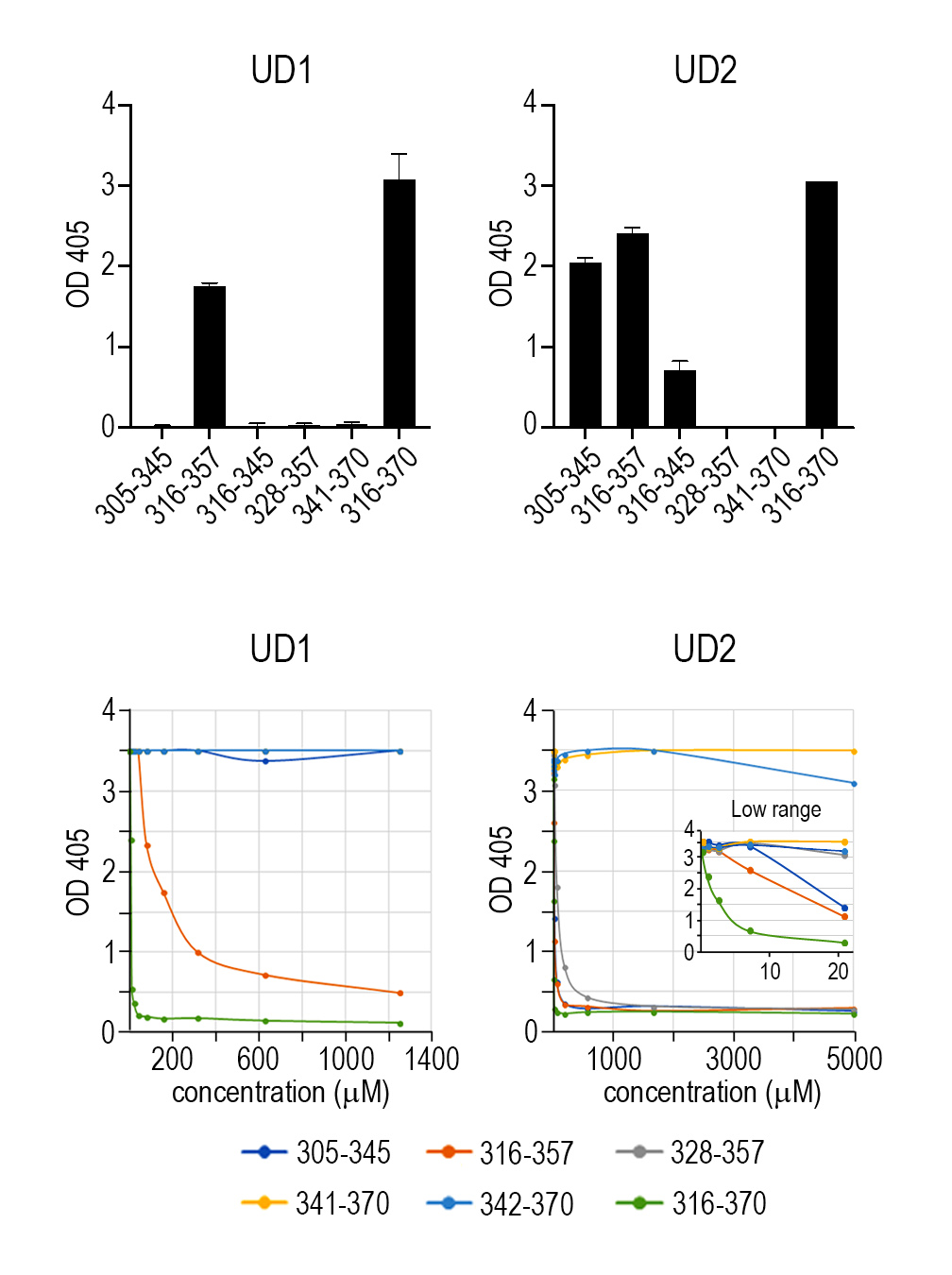
