## Supplement Description for "Uman Type NF-L Antibodies Are Effective Reagents for the Imaging of Neurodegeneration"

**Supplementary Material**

**Supplement figure 1: Peptide inhibition assays.** Mapping of epitopes of UD2 and other NF-L antibodies by inhibition of antibody binding to human NF-L by 20 amino acid nested peptides overlapping by 5 amino acids. These experiments provided no information for several NF-L antibodies including notably UD1 and MCA-1B11.

**Supplement figure 2:** **Direct peptide binding and competition experiments.** The results of testing UD1 and UD2 binding to NF-L peptides 305-345, 316-357, 316-345, 328-357 and 341-370 applied in triplicate and equimolar amounts to ELISA plates. **A:** UD1 binds 316-370 and less well to 316-357. **B:** In the same assay UD2 binds 316-370 and less well to 305-345, 316-357 and 316-345, focusing attention on the 316-335 in line with other data. **C, D** We challenged the binding of UD1 and UD2 to full length recombinant human NF-L in an ELISA format with a range of equimolar concentrations of the 5 peptides. In both cases and in line with direct binding data the 316-370 was a very efficient inhibitor of antibody binding to NF-L. **C:** For UD1 316-357 showed less efficient inhibition at the range of concentrations tested. **D:** In the case of UD2 all peptides containing the 316-335 showed significant but less potent inhibition. Inset shows inhibition at very low peptide concentrations.
