## Supplement Table 1 for "Uman Type NF-L Antibodies Are Effective Reagents for the Imaging of Neurodegeneration"

**Supplementary Table 1: Antibodies**

| **Name and link** | **Source** | | | | | **Type*** | | | **Immunogen** | | | **Specificity** | **Reference** | |
| --- | --- | --- | --- | --- | --- | --- | --- | --- | --- | --- | --- | --- | --- | --- |
| [UD1/2.1](https://www.umandiagnostics.se/cms/neurofilament-light-antibodies/) | | Uman | | | M-MA | | | HPLC purified mammalian spinal cord NF-L | | | Human NF-L 322-357 | | | 1 |
| [UD2/47.3](https://www.umandiagnostics.se/cms/neurofilament-light-antibodies/) | | Uman | | | M-MA | | | HPLC purified mammalian spinal cord NF-L | | | Human NF-L 316-330 | | | 1 |
| [MCA-DA2](https://encorbio.com/product/mca-da2) | | EnCor | | | M-MA | | | Dephosphorylated pig neurofilament preparation | | | Human NF-L 441-455 | | | 2 |
| [MCA-6H63](https://encorbio.com/product/mca-6h63) | | EnCor | | | M-MA | | | Proprietary immunogen including NF-L 311-362 | | | Human NF-L 311-315 | | | NA |
| [MCA-1D44](https://encorbio.com/product/mca-id44) | | EnCor | | | M-MA | | | Proprietary immunogen including NF-L 311-362 | | | Human NF-L 316-330 | | | NA |
| [MCA-1B11](https://encorbio.com/product/mca-1b11) | | EnCor | | | M-MA | | | Purified pig spinal cord NF-L | | | Human NF-L 322-357 | | | NA |
| [MCA-3H11](https://encorbio.com/product/mca-3h11) | | EnCor | | | M-MA | | | Recombinant rat NF-M tail | | | Rat NF-M tail, 762-845 | | | 2 |
| [MCA-2E3](https://encorbio.com/product/mca-2e3) | | EnCor | | | M-MA | | | Recombinant rat α-internexin | | | Human α-internexin 340-505 | | | 3 |
| [MCA-1B63](https://encorbio.com/product/mca-1b63) | | EnCor | | | M-MA | | | Recombinant pET30a(+) leader sequence | | | S-tag peptide | | | NA |
| [MCA-NAP4](https://encorbio.com/product/mca-nap4) | | EnCor | | | M-MA | | | Axonal NF-H purified from bovine spinal cord | | | NF-H phospho-KSP sequences | | | 4 |
| [MCA-5B10](https://encorbio.com/product/mca-5b10) | | EnCor | | | M-MA | | | Recombinant human low molecular weight 3 repeat MAPτ | | | Tau isoform hTau40 362-381 | | | 5 |
| [RPCA-NF-L-ct](https://encorbio.com/product/rpca-nf-l-ct) | | EnCor | | | R-PA | | | C-terminal 29 amino acids of rat NF-L | | | Rat NF-L C-terminus 515-543 | | | NA |
| [RPCA-NF-L-Degen](https://encorbio.com/product/rpca-nf-l-degen) | | EnCor | | | R-PA | | | Recombinant human NF-L 311-362 | | | Human NF-L 311-362 | | | NA |
| [CPCA-NF-L-Degen](https://encorbio.com/product/cpca-nf-l-degen) | | EnCor | | | C-PA | | | Recombinant human NF-L 311-362 | | | Human NF-L 311-362 | | | NA |
| [Anti-IFA/TIB-131](https://www.atcc.org/products/tib-131) | | ATCC | | | M-MA | | | GFAP from post mortem human spinal cord | | | Human NF-L 381-400 | | | 6 |
| [Ab18184](https://www.abcam.com/6x-his-tag-antibody-hish8-ab18184.html) | | Abcam | | | M-MA | | | Poly histidine peptide | | | 6 sequential histidine residues | | | NA |
| [4F8](https://pubmed.ncbi.nlm.nih.gov/27503460/) | | NA | | | M-MA | | | NF-L peptide 465-480 including phospho Ser-473 | | | Phospho-Ser 473 NF-L 465-480 | | | 7 |

*M-MA = mouse monoclonal antibody, R-PA = affinity purified rabbit polyclonal antibody, C-PA = affinity purified chicken polyclonal antibody
